# miRNA Mimic Optimization: Chemical Structure – Activity – Targetome Relationship to Engineer Selective Anti-Tumor Immunity in T cells

**DOI:** 10.1101/2025.04.16.649059

**Authors:** Xavier Segarra-Visent, Tatyana Ryaykenen, Anastasia Kremer, Melanie Sauer, Samuel Jackson, David A Cooper, Dimas Echeverria, Reka A Haraszti

**Affiliations:** Department of Internal Medicine II, Hematology, Oncology, Clinical Immunology and Rheumatology, University Hospital Tuebingen; Gene and RNA Therapy Center (GRTC), Faculty of Medicine, University Tuebingen; RNA Therapeutics Institute, University of Massachusetts Chan School of Medicine, Worcester, MA 01605, USA

## Abstract

MicroRNA (miRNA) mimic therapies act through a broad and complex targetome in disease contexts. However, both the composition of these targetomes and the impact of chemical modifications on them remain poorly understood.

In this study, we investigate eight fully chemically modified miRNA scaffolds across three miRNA sequences in a model immune disorder, graft-versus-host-disease (GvHD), which is an off-tumor effect of allogeneic T cell therapy. We demonstrate that conventional silencing assays fail to predict the functional performance of miRNA mimics in GVHD and graft-versus-leukemia (GvL), the on-tumor effect of allogeneic T cells. Moreover, we find that chemical scaffolds influence the duration of silencing mediated by miRNA mimics.

We identify a miR-374b version as a lead miRNA candidate that not only inhibits GVHD but also enhances GVL—an unprecedented and desired improvement in on-versus-off-tumor selectivity of clinical relevance. Further analysis reveals distinct responder and non-responder groups to miR-374b therapy. While responders exhibit significant transcriptomic shifts, non-responders show virtually no changes, despite intact miRNA pathway function, underscoring a strong correlation between transcriptome reprogramming and therapeutic efficacy. RNA sequencing reveals that miR-374b acts via metabolic reprogramming of T cells and induces partially distinct transcriptional programs in GVHD and GVL.

Collectively, our findings demonstrate that fully chemically modified miRNA mimic therapies are feasible to rewrite the T-cell transcriptome, profoundly improving on-versus-off-tumor specificity in T-cell-mediated cancer therapies.

## Introduction

miRNAs are small inhibitory RNAs that act as master regulators of homeostasis^1,2^, and their dysregulation is a common hallmark of various diseases^3,4^. Therefore, the supplementation of miRNAs *via* miRNA mimics is an important therapeutic goal.

Initial enthusiasm for miRNA-based therapies was dampened by a series of failed early clinical trials that suffered from immune-mediated adverse events outweighing low levels of efficacy^5,6^ – similar to early siRNA trials^7^. Using unmodified^5,6^ or partially chemically modified^8^ miRNA mimics likely substantially contributed to these failures. Lessons from the siRNA field suggest that chemical modifications are likely necessary to harness the therapeutic potential of miRNA mimics^9^. The siRNA field discovered that full chemical modification is required for a biologically meaningful effect size and duration *in vivo*^10^, and that such modifications can also eliminate the immune activation intrinsic to small double-stranded RNAs^11,12^. However, applying this knowledge to miRNAs has been challenging due to a key difference: miRNAs tolerate mismatches when targeting mRNAs for degradation, resulting in an unknown, bioinformatically not entirely predictable and potentially large number of mRNA targets (targetome)^3^. This uncertainty makes it difficult to precisely identify on-target and off-target effects of miRNA-based therapies, complicating the development of miRNA mimics^13^. Chemical modifications can influence the melting temperature of oligonucleotide-RNA duplexes, especially at mismatch sites^14^. This feature has been used in siRNAs to create allele-specific compounds^14^, but in miRNAs, it rather increases unpredictability. Introducing a miRNA mimic to cells affects a large and unknown number of targets, and chemical modifications can differently influence subsets of these targets^9^. There have been sporadic reports^15–19^ of fully chemically modified miRNAs showing functional efficacy in mouse models^15–17,19^, yet, characterization of the miRNA targetome mediating *in vivo* efficacy is partially lacking^15,17,19^. miRNA mimics used in these studies incorporate an alternating 2’-O-methyl and 2’-fluoro modification pattern^15,16,19^, with either a full-length passenger strand containing mismatches^15,19^ or a fully matched but shortened passenger strand^16^ sequence. Chemical optimization of miRNA mimics used in above mouse studies has not been reported^15–19^. Recently, extensive *in vitro* chemical optimization of miRNA mimics has revealed that 1) full chemical modification impairs miRNA activity compared to partially modified or unmodified versions, 2) fully matched, full-length passenger strands reduce silencing efficacy, and 3) different chemical modification patterns affect silencing activity on different targets to a different extent^9^. However, no functional assays in disease models were conducted^9^. As a result, it remains unclear how chemical optimization of miRNA mimics impacts their biological function in diseases and how these effects are mediated through the miRNA targetome.

In the present study we chemically optimize fully modified miRNA mimics in a model immune disorder— graft-versus-host-disease (GvHD), an immune-related complication of allogeneic hematopoietic stem cell transplantation (HSCT). HSCT is a curative cell therapy for many types of leukemia and lymphoma, where transplanted allogeneic T cells recognize and eliminate leukemia cells through the graft-versus-leukemia effect (GvL) effect. However, these same T cells can also attack healthy tissues, causing GvHD as an off-tumor toxicity. GvHD and GvL co-occur in 15–26%^20,21^ of cases, while GvHD can appear independently of GvL in 5–34%^21,22^ of cases, and GvL can manifest without GvHD in 24–46%^22,23^ (these statistics take into account the routine use of GvHD prophylaxis). Thus, although GvHD and GvL correlate^24^, GvHD-free, relapse-free survival^22,23^ is evidence that leukemia-specific immunity can be achieved in the context of HSCT.

Several miRNAs^25^, including miR-146a^26^, miR-181a^27^, and miR-374b^28,29^, have been found to be dysregulated in GvHD in T cells. T-cell-specific knockouts of miR-146a and miR-181a loci aggravated GvHD in mice^26,27^. Single nucleotide polymorphisms in miR-146a loci correlated with GvHD severity in patients^26^. Furthermore, miRNA replacement with either commercially available partially chemically modified miR-146a mimic^26^ or lentivirally expressed miR-181a^27^ have demonstrated beneficial effects in mitigating GvHD in mice. Downregulation of TRAF6 by miR-146a^26^ and BLC2 by miR-181a^27^ was suggested to mediate these effects in GvHD. Importantly, miR-146a and miR-181a replacement was found to preserve GvL effects, while inhibiting GvHD^25^ –addressing a major precision medicine challenge in immune-oncology: enhancing antitumor responses without exacerbating auto/alloimmune complications. miR-374b^30^ and miR-146a^31^ have been also shown to be upregulated in regulatory T cells, which are known to alleviate GvHD^32^. However, the mechanistic role of miR-374b in GvHD remains largely unexplored. While the abundance of miRNAs in T cells varies depending on their developmental and functional state, miR-181a (260–10,000 TPM, constituting ∼13% of all miRNAs in lymphocytes) and miR-146a (10– 2,800 TPM) rank among the most abundant miRNAs in T cells^33^. miR-374b is expressed at lower levels, typically between 30 and 560 TPM^33^. Comprehensive targetome analyses for these miRNAs in T cells are still lacking, leaving room to further elucidate the mechanisms underlying functional role in GvHD/GvL. Beyond investigating the mechanistic roles of above miRNAs in GvHD, the lack of optimized chemical modifications for miRNA mimics poses a further significant challenge to advancing miRNA replacement therapies for this condition.

Here, we aim to fill this gap by systematically screening full chemical modification patterns of miR-146a, miR-181a, and miR-374b mimics in silencing assays and GvHD/GvL models.

## Results

We used a fully chemically modified miRNA mimic pattern featuring alternating 2’-fluoro and 2’-O-methyl groups, as commonly reported in all *in vivo* studies using fully modified miRNA mimics^15–17^, with 2’-O-methyl at the first nucleotide of the guide strand’s 5’ end (Figure 1). In scaffolds M1 and M2, we used both guide and passenger strand sequences from miRBase^34^ for mature human miRNAs miR-146a, miR-181a and miR-374b. For scaffolds M3 and M4, we modified passenger strands to remove mismatches and make them fully complementary to the respective guide strands (fully matched, Figure 1).

**Figure 1.**
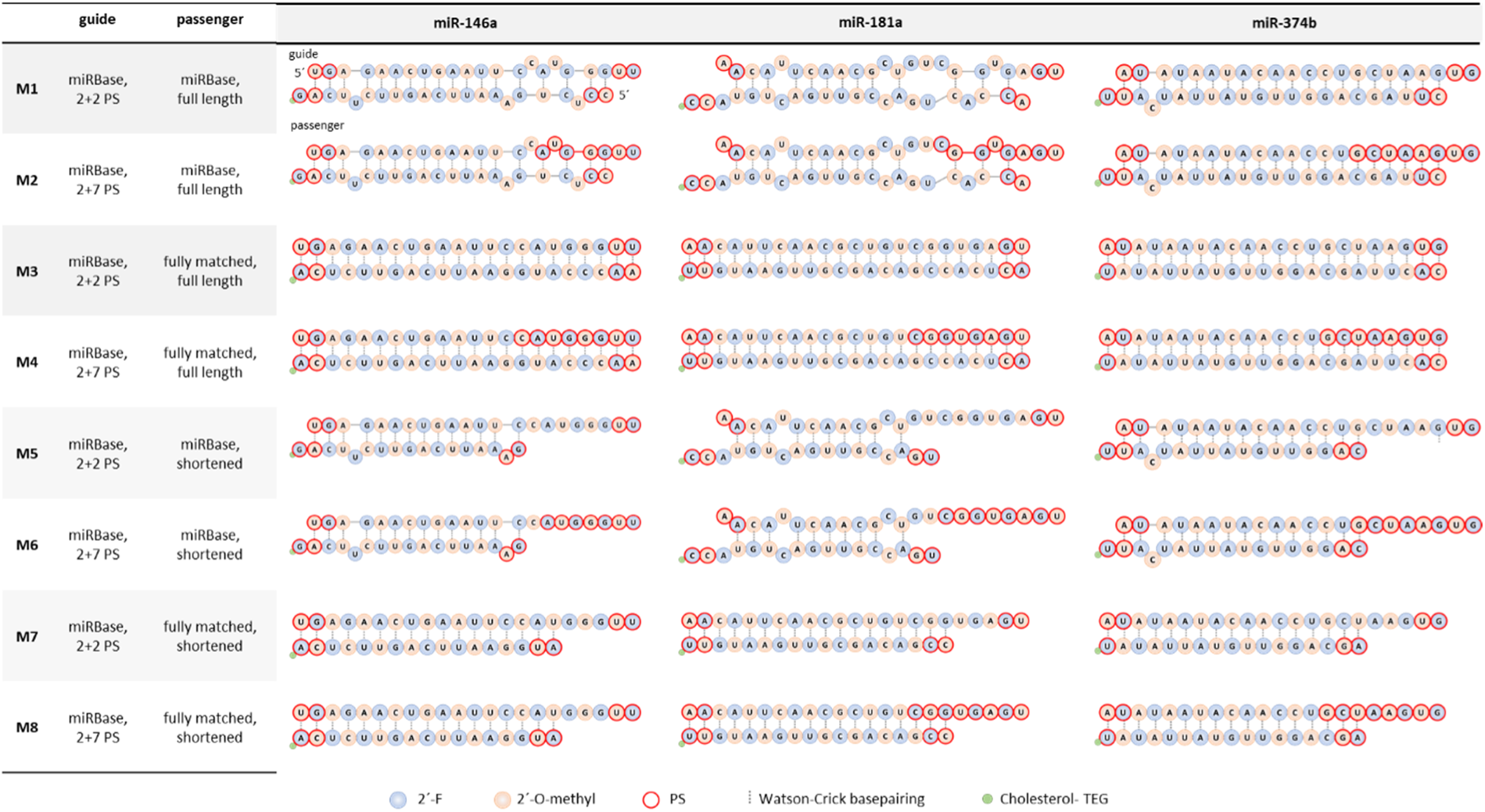
Chemical structures of miRNA mimics. The sequences of human mature miR-146a, miR-181a and miR-374b were obtained from miRbase^34^. miRNAs were modified with an alternating pattern of 2’-O-methyl (orange) and 2’-Fluoro (blue), with 2’-O-methyl at the 5’-end of the guide strand. 3’-ends of the passenger strands were covalently conjugated to cholesterol with a TEG linker (green). Phosphorothioate modifications are depicted in red. Watson-Crick base pairing is shown as a dashed line.

To examine the impact of duplex region length, we synthesized truncated passenger strands (shortened by 5 nucleotides) using either the respective miRBase sequence (M5 and M6) or sequences fully matched to the guide (M7 and M8). We stabilized both ends of passenger strands and the 5’ end of guide strands with two phosphorothioate (PS) linkages in all scaffolds. The 3’ end of guide strands was stabilized with either two (M1, M3, M5, and M7) or seven PS linkages (M2, M4, M6, and M8). Additionally, all guide strands were phosphorylated at the 5’ end, and all passenger strands were covalently conjugated to cholesterol *via* a triethyl glycerol (TEG) linker at the 3’ end.

We synthesized miRNA mimic chemical scaffolds M1 to M8 for three miRNAs indicated in GvHD biology, each exhibiting different degrees of complementarity between their strands: miR-374b with 1 mismatch and 20 Watson-Crick base pairs, miR-146a with 5 mismatches and 18 base pairs, and miR-181a with 7 mismatches and 13 base pairs (Figure 1).

### Shortening fully matched passenger strands rescues miRNA mimic silencing impairment on fully complementary reporter target

We first validated the fully chemically modified miRNA mimics in a luciferase reporter assay, where we cloned target sequences fully complementary to the miRNA guide sequence (four times each) into the 3’UTR of Renilla Luciferase in the psiCHECK2 plasmid (Fig.2A). In general, we observed some level of silencing activity across all fully chemically modified miRNA mimics. Full sequence complementarity between strands (M3 and M4) appeared to slightly impair activity compared to M1 and M2 for miR-181a and miR-146a but not for miR-374b, whose natural sequence contained only one mismatch (Fig.2A). Truncating the passenger strand had little effect on activity of duplexes with the natural passenger strand sequence (M5 and M6 versus M1 and M2), but it partially restored the lost activity in miRNAs containing completely matched passenger strands (M7 and M8 versus M3 and M4) (Fig.2A). In asymmetric duplexes, where the guide strand has a 5 nucleotides long single-stranded overhang, we expected increased number of PS linkages in this overhang region to enhance activity. We observed this advantage of extended PS linkages (M7 versus M8 and M5 versus M6) across all three miRNA mimics, except in the context of mismatched natural passenger strands of miR-181a (M5 versus M6), where additional PS linkages unexpectedly reduced activity (Fig.2A). Interestingly, additional PS linkages also enhanced silencing activity of miRNA mimics with full length passenger strands (M1 versus M2) for miR-181a and miR-146a in the context of natural mismatched passenger strands, but, conversely, reduced activity for miR-374b (Fig.2A).

**Figure 2.**
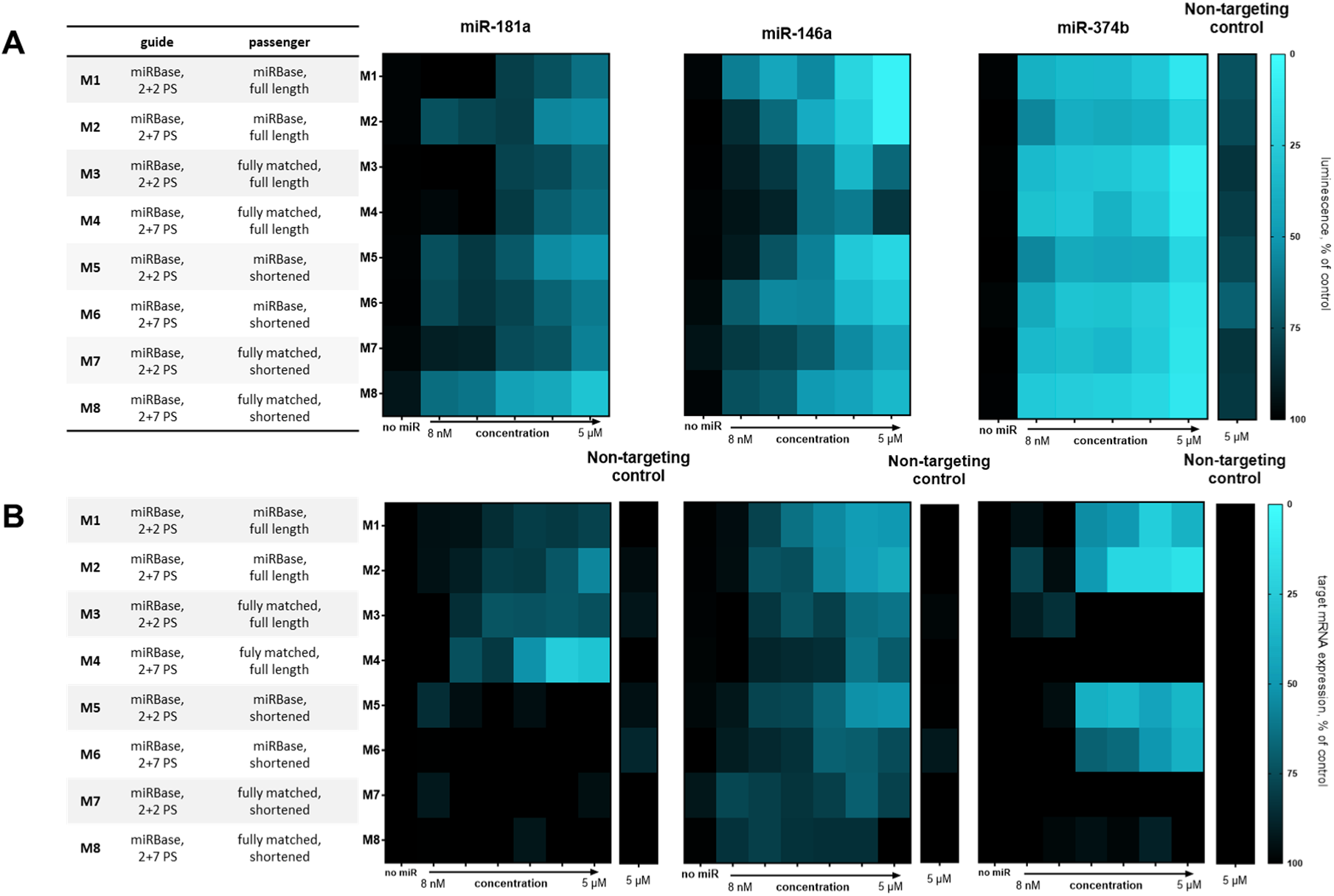
miRNA mimic silencing activity on reporter and natural targets. (A) HeLa cells with a reporter plasmid including fully complementary target sites for the miRNAs were treated with various concentrations of miRNA mimics and non-targeting controls, as shown on the x axis. After 3 days of co-incubation, firefly and Renilla luminescence were analyzed with a DualGlo reporter system, and expression of the reporter was calculated by dividing the Renilla signal (on-target), with the firefly signal (transfection control) and normalized to untreated samples. Reporter expression was color-coded with black representing no silencing and blue representing maximal silencing. N=3-6 (B) Jurkat cells were treated with miRNA mimics at various concentrations indicated on the x axis. Silencing of previously identified miRNA targets (*AKT1* – miR-181a, *IRAK1 –* miR-146a and *PTPN11* – miR-374b) was quantitated after 6 days of incubation using QuantiGene SinglePlex assay and *HPRT* as housekeeping gene. Target gene expression was normalized to untreated samples and color-coded with black representing no silencing and blue representing maximal silencing. (N=3-8)

### Shortened fully matched passenger strands abolish miRNA mimic silencing activity on natural targets

We then sought to evaluate the activity of fully chemically modified miRNA mimics on validated natural targets. For this, we selected PTPN11^35^, IRAK1^26^ and AKT1^36^ as experimentally validated targets for miR-181a, miR-146a and miR-374b, with 4, 3 and 2 predicted binding sites (according to miRanda), respectively. First, we generally observed reduced silencing activity (Figure 2B) compared to the fully complementary reporter target (Figure 2A), with a notably different pattern in terms of which chemical scaffolds were active or inactive. Strikingly, truncating the passenger strand completely abolished miR-181a activity on PTPN11 (M5-M8 versus M1-M4), whereas a fully matched passenger strand significantly improved activity, particularly with extended PS linkages (M4 versus M2) (Fig.2B). In contrast, for miR-146a and miR-374b, the fully matched passenger strand diminished activity on IRAK1 and AKT1 (M3-M4 versus M1-M2), especially when shortened (M7-M8 versus M5-M6) (Fig.2B). miR-146a mimic containing fully matched shortened passenger strands remained functional, with no substantial effect observed from PS linkages (M7-M8) (Fig.2B).

We next asked whether differences in cellular uptake or endosomal escape of the various chemical scaffolds could account for the observed variations in silencing activity. To explore this, we transfected miR-146a scaffolds M1-M8 into cells using RNAiMax and compared their silencing activity to that of miR-146a mimics introduced *via* passive uptake (Suppl. Fig.1A). The resulting patterns were very similar, suggesting that interactions with AGO2 and target mRNA, rather than cellular uptake mechanisms, are responsible for the variations in activity of fully chemically modified miRNA mimics. A further question was whether chemical modification patterns would affect duration of silencing and this could be a basis of selecting one chemical architecture for downstream applications. To address this, we used cells stably transduced with a fluorescent miRNA reporter mimicking a natural target with central mismatch bulk^37^ (Supp.Fig.1B). At early timepoints we saw similar variance in silencing activity than previously observed with passive uptake with M1, M2, M3 and M5 structures being most active (Supp. Fig.1B). Then we saw a steep drop in activity with structures M2 and M5 at approximately one-week post-treatment, while M1 and M3 maintained silencing beyond three weeks (Supp. Fig.1B). Generally, structures with higher PS content and/or shortened passenger strands exhibited shorter silencing durations.

Overall, we observed that chemical modification patterns and miRNA sequence work together to determine silencing activity. Yet, previous work has shown that silencing activity of chemically modified miRNA mimics can substantially differ on distinct mRNA targets^9^. A crucial unanswered question remains whether and how experimentally observed variations in silencing activity of miRNA mimics can reliably predict functional outcomes in disease contexts. This relationship has not been previously investigated, leaving a significant gap in understanding the translational requirements of chemically modified miRNA mimic therapies.

### miRNA mimic PS content is the strongest predictor of activity in in vitro GvHD models

To address this gap, we proceeded to evaluate chemically modified miRNA mimics in an *in vitro* GvHD assay (Fig.3A). All miRNA mimics demonstrated significant activity in inhibiting T cell proliferation in response to allogeneic stimulation, with those containing higher PS content generally showing superior activity (Fig.3A). This enhanced performance may be due to increased non-specific protein binding mediated by PS^38^. Interestingly, even scaffolds that were ineffective at silencing the tested natural targets (such as M5-M8 of miR-181a and M3-M4/M7-M8 of miR-374b, Fig.2B) were still able to inhibit T cell proliferation in an *in vitro* GvHD assay (Fig.3A), likely due to retained activity on other natural targets relevant to GvHD. It has been previously shown that certain chemical modifications can abolish silencing on one target while preserving efficacy on another^9^. Importantly, the observed effect was sequence-specific, as non-targeting controls did not inhibit T cell proliferation (Fig.3A). Notably, the GvHD inhibitory effect of the tested miRNA mimics was comparable to that of approved GvHD drugs (Fig.3A). We next asked whether the inhibitory effect of miRNA mimics was specific to GvHD. To address this, we first tested the impact of miRNA mimics on polyclonally activated T cells (*via* CD2/CD3-CD28 beads), which are not expected to be inhibited by a GvHD-specific therapy. We observed no inhibition of polyclonally activated T cells by miRNA mimics, while most approved GvHD drugs led to substantial inhibition (Fig.3B). Yet, polyclonal activation depicts T cell activity in an artificial context. More relevant to GvHD-specific therapies would be the ability of T cells to attack tumor cells, *i.e.* the GvL effect.

**Figure 3.**
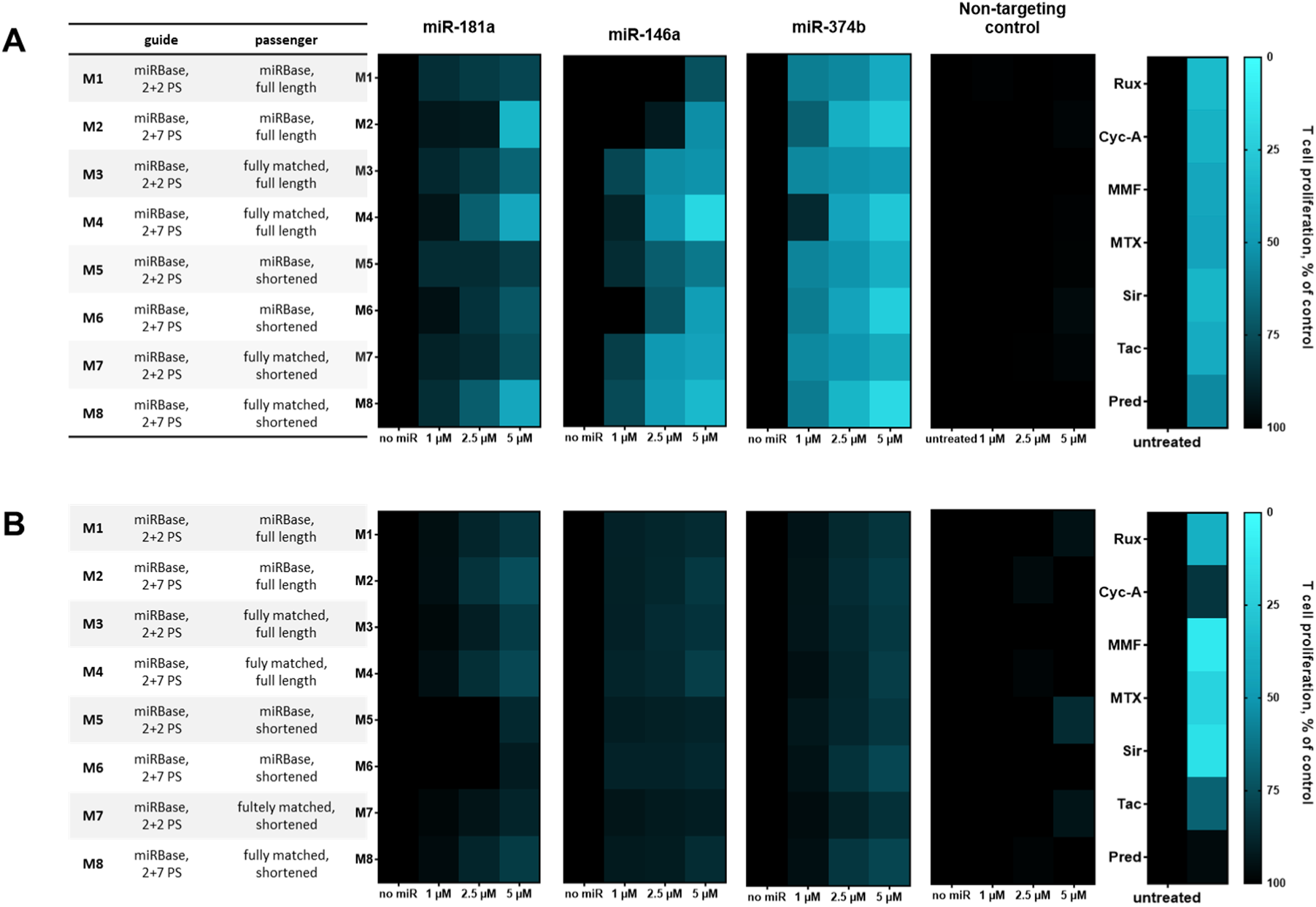
miRNA mimic functional activity in a GvHD model and in polyclonally activated T cells. Responder T cells were co-incubated with allogeneic irradiated PBMCs (A) or CD2/CD3/CD28 activation beads (B). Co-cultures were treated with miRNA mimics or approved GvHD drugs (Rux: 7.5 µM Ruxolitinib, Cyc-A: 5 µM Cyclosporin A, MMF: 10 µM Mycophenolat-Mofetil, MTX: 100 µM Methotrexate, Sir: 25 µM Rapamycin, Tac: 10 µM Tacrolimus, Pred: 7.5 µM Prednisolone) at various concentrations as depicted on the x axis, and incubated for 5 days. T cell proliferation was then quantified based on dilution of the CellTraceViolet proliferation dye and flow cytometry. T cell proliferation data were normalized to untreated co-cultures and color coded with black depicting no reduction in T cell proliferation, and blue depicting maximal reduction of T cell proliferation. N=3

### miR mimic chemical scaffold affects GvL

We therefore aimed to assess whether miRNA mimics compromise the GvL effect, which would limit their potential as a precision therapy for GvHD. In an *in vitro* GvL assay, we observed that miRNA mimics either preserved or reduced leukemia viability (Fig.4A), suggesting that they preserve or even enhance the GvL effect. This enhancement was sequence-specific, as non-targeting miRNA mimics had no impact on leukemia cell viability in GvL co-cultures (Fig.4A). In contrast, approved GvHD drugs led to increased leukemia cell numbers and viability in GvL assays (Fig.4A), indicating a weakened GvL effect. This aligns with clinical observations of leukemia relapse following intensive GvHD treatment. Notably, the reduced GvL effect induced by approved drugs was associated with suppressed T cell proliferation in GvL co-cultures (Fig.4B), leading to up to a tenfold decrease in T cell numbers. miRNA mimics, however, either had no effect or induced only a minor reduction in T cell numbers (Fig.4B) in GvL assays.

**Figure 4.**
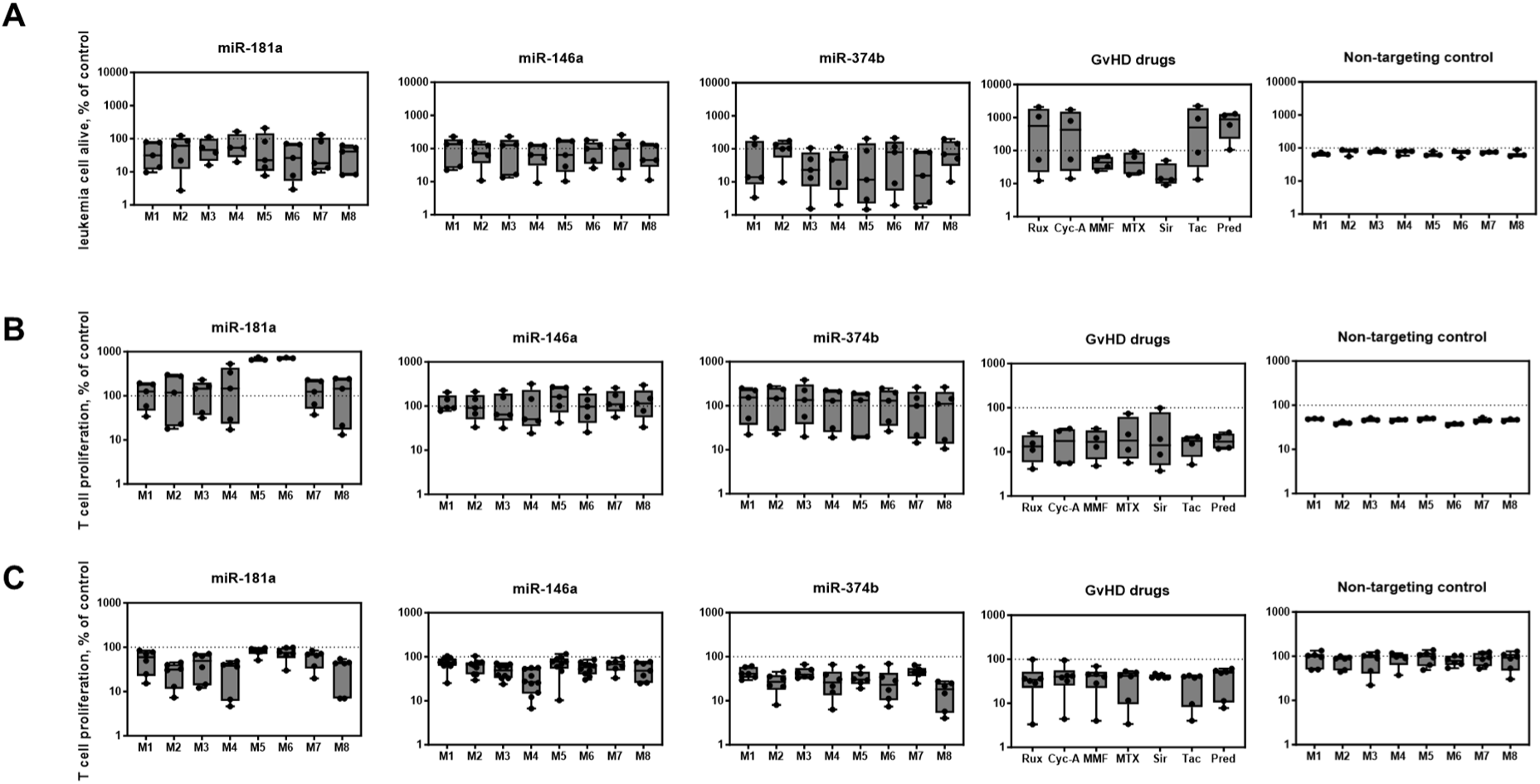
miRNA mimics functional activity in GvL and GvHD model. Responder T cells were co-incubated with allogeneic, irradiated dendritic cells (A-C) and in some cases also with the MOLM-13 acute myeloid leukemia cell line (A-B). Co-cultures were treated with miRNA mimics at 5µM or with GvHD drugs (Rux: 7.5 µM Ruxolitinib, Cyc-A: 5 µM Cyclosporin A, MMF: 10 µM Mycophenolat-Mofetil, MTX: 100 µM Methotrexate, Sir: 25 µM Rapamycin, Tac: 10 µM Tacrolimus, Pred: 7.5 µM Prednisolone) for 5 days. T cell proliferation (B-C) and viable MOLM-13 cells (A) were then quantified using flow cytometry. Each dot represents and individual experiment. Box plot showing minimum and maximum (whisker), 95% confidence interval (box) and mean (line). N=3-10

When comparing T cell proliferation inhibition in GvL versus in GvHD, we found that approved drugs suppressed T cells more potently in GvL assays (cca. 10-fold, Fig.4B) than in GvHD assays (cca. 3-fold, Fig.4C), p=0.004. In contrast, miRNA mimics exhibited the opposite pattern, inhibiting T cell proliferation more in GvHD assays (Fig.4C) than in GvL assays (Fig.4B) (no inhibition in GvL vs. up to 10-fold inhibition in GvHD, p<0.0001 with all miRNA sequences tested). In some cases, miRNA mimic treatment even enhanced T cell proliferation in GvL assays (Fig.4B).

While response variability was broad, we found no consistent correlation between GvL outcomes and the chemical modification scaffolds of miRNA mimics (Fig.4A and 4B). However, these modification scaffolds significantly influenced miRNA mimic-mediated inhibition of T cell proliferation in GvHD co-cultures (Fig.4C).

The broad range of responses observed in these *in vitro* assays likely reflects the inherent batch-to-batch variability of T cells, which is also observed in clinical patient responses. Collectively, our findings suggest that miRNA mimics represent a promising therapeutic strategy for inducing tumor-selective immune responses in the allogeneic setting. However, the specific pathways mediating their effects in GvHD and GvL remain unclear.

### GvHD and GvL responders to miR-374b M4 overlap

Next, we aimed to characterize the targetome of a chemically modified miRNA mimic in the context of GvHD and GvL. We selected the M4 scaffold of miR-374b for this analysis, as it demonstrated the strongest dual effect—maximizing GvHD inhibition while enhancing the GvL response (Fig.3 and 4). Notably, we observed a broad range of responses to the M4 miR-374b mimic in both GvHD and GvL models (Fig.4), with populations of responder and non-responder T cells. This variability mirrors what is commonly seen in T-cell-mediated cancer therapies in the clinic, where patients often show diverse responses, categorized as responders or non-responders. Consequently, we also sought to determine whether the targetome of miR-374b differs between responder and non-responder T cells, potentially shedding light on the mechanisms driving this differential efficacy.

We first aimed to determine whether T cells responsive to the miR-374b M4 mimic in the GvHD assay are the same cells that also exhibit a response in the GvL assay. To address this, we conducted paired experiments using T cells from the same donor for both assays (Supp. Fig.2). Our results revealed a correlation: T cells that demonstrated greater GvHD inhibition also showed enhanced GvL activity in response to the miR-374b M4 mimic (Supp. Fig.2.). We selected subsets of T cells from these experiments: GvHD non-responders, which were GvL non-enhancers at the same time (Supp. Fig.2, depicted in blue) and responders for both GvHD inhibition and GvL enhancement (Supp. Fig.2, depicted in orange). To uncover the underlying mechanisms, we performed bulk RNA sequencing (RNA-seq) on these distinct T-cell populations, aiming to define mechanisms affecting the response to miR374b M4 mimic in GvHD and GvL co-culture contexts.

### The scope of the regulated targetome defines therapeutic response to miR-374b-mimic

We identified 697 differentially expressed genes (DEGs) in miR-374b-M4-treated *versus* untreated GvHD responder T cells (Fig.5A) and 1,084 DEGs in miR-374b-M4-treated *versus* untreated GvL responder T cells (Fig.5B). Interestingly, the number of DEGs in GvHD non-responders and GvL non-responders upon miR-374b-M4 treatment was substantially lower (42 and 49, respectively) (Fig.5C and 5D). These findings suggest that the number of DEGs correlate strongly with the effect size in functional GvHD and GvL assays.

**Figure 5.**
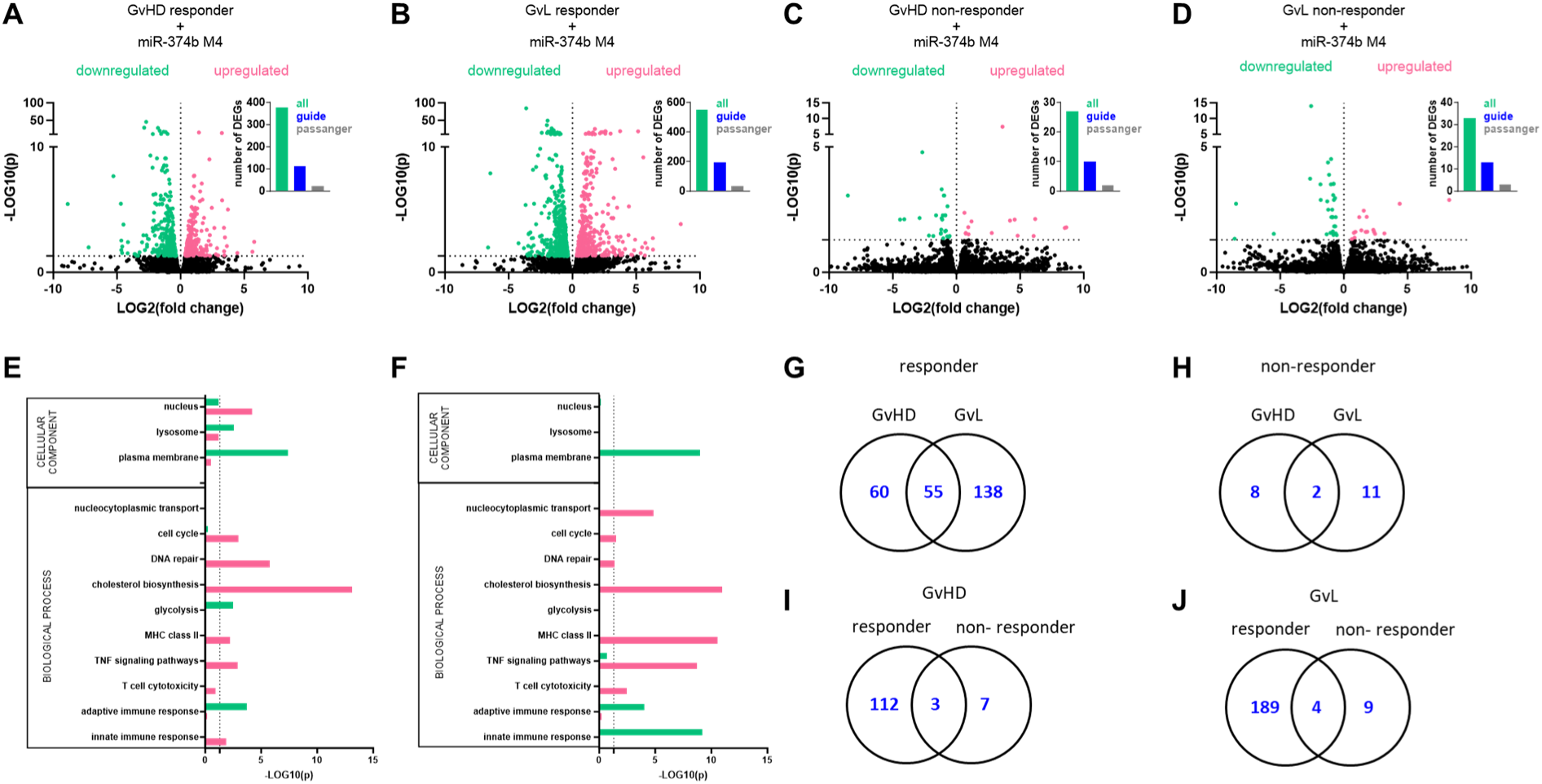
miR-374b-M4 targetome in responder and non-responder GvHD and GvL models. T cells were re-purified from responder and non-responder co-cultures of GvHD and GvL models, RNA purified and subjected to bulk RNASeq. Differentially expressed genes (DEGs, paired analysis) between miR-374b-M4-treated and untreated cells are shown in GvHD responder T cells (A), GvL responder T cells (B), GvHD non-responder T cells (C) and GvL non-responder T cells (D). Downregulated genes are depicted in green and upregulated genes in magenta. Downregulated genes were further analyzed for containing seed matches in the 3’-UTRs (A-D, inlets) for the miR-374 guide strand (in blue) or passenger strand (in grey). Gene ontology analysis was performed on DEGs obtained from responder GvHD (E) and responder GvL (F) co-cultures. Venn diagrams (G-J) show overlaps in downregulated genes with a seed match to miR-374b guide strand (in blue) in responder GvHD *versus* GvL cells (G), in non-responder GvHD *versus* GvL cells (H), in responder *versus* non-responder GvHD cells (I) and in responder *versus* non-responder GvL cells (J). N=2

In both responders and non-responders, about one-third of all downregulated DEGs contained a seed match to the miR-374b guide strand, while only 6-7% contained a seed match to the miR-374b passenger strand. This suggests two key points: (1) The M4 chemical modification platform ensures correct strand selection by AGO2, and (2) miR-374b’s effects in responders are potentially mediated *via* hundreds of on-targets, making it challenging to identify which of these targets are truly relevant to the disease mechanisms.

To further explore the mechanisms behind miR-374b-M4’s action in responders, we first performed gene ontology (GO) analysis on all DEGs (Fig.5E and 5F). In both GvHD and GvL, we observed significant downregulation of genes encoding plasma membrane proteins upon miR-374b-M4 treatment (Fig.5E and 5F). In GvHD, there was also notable regulation of lysosomal and nuclear proteins (Fig.5E), which was not seen in GvL (Fig.5F). Consistent with this, cell cycle and DNA-repair-related genes were more significantly regulated in GvHD (Fig.5E) than in GvL (Fig.5F), which aligns with our previous observation (Fig.4) that miR-374b-M4 only inhibits T cell proliferation in GvHD, but not in GvL. Interestingly, miR-374b-M4 upregulated nucleocytoplasmic transport in GvL responders, suggesting substantial changes in transcriptome regulation, as many transcription factors in T cells shuttle between the cytoplasm and nucleus.

In terms of biological processes, the most prominent effect was the upregulation of cholesterol biosynthesis in both GvHD and GvL responders upon miR-374b-M4 treatment (Fig.5E and 5F). Conversely, we observed downregulation of the glycolysis pathway only in GvHD (Fig.5E) but not in GvL (Fig.5F). These data are consistent with previous literature highlighting glycolysis as a target pathway in GvHD^39–41^ and the role of lipid metabolism in T cell activation and allogeneic immune responses.

miR-374b-M4 treatment in responders also modulated immune regulatory pathways. Both GvHD and GvL responder T cells showed upregulation of MHC class II and TNF signaling pathways, along with downregulation of adaptive immune responses upon treatment with miR-374b-M4. MHC class II, typically expressed on antigen-presenting cells but also a late activation marker on T cells^42^. TNF-α, a pro-inflammatory cytokine, is associated with anti-tumor T cell responses^43^. Therefore, we speculate that upregulation of MHC-II and TNF-α likely contributes to GvL. Their role in inhibiting GvHD, however, is unclear. In contrast, the downregulation of the adaptive immune response may help in reducing GvHD but seems disadvantageous in GvL.

The regulation of the innate immune response was a major differentiator between GvHD and GvL responder T cells: it was upregulated in GvHD responders and downregulated in GvL responders after miR-374b-M4 treatment. The context is that GvL responders showed much higher expression of interferon-induced genes compared to GvHD responders. We didn’t see this effect in non-responders, indicating that responder T cells may be particularly sensitive to type I interferon responses, which are attenuated by miR-374b-M4 in GvL and enhanced in GvHD. Type I interferons have context-dependent effects on T cells^44^. In some models, type I interferons protect from inflammatory responses in the gut, including in GvHD^45,46^. Type I interferons are known to exacerbate on-tumor immune responses in various contexts, including in GvL^45,47^. However, type I interferons also induce T cell exhaustion markers such as PD-1, TIM-3, and LAG-3^48^, suggesting that curtailing the overactivation of type I interferon signaling in GvL may be beneficial.

We then went on to examine the putative on-targets (downregulated gene with seed match to the miR guide strand) of miR-374b-M4 treatment. In responders, 55 miR targets were shared between GvHD and GvL, whereas a substantial number of targets were distinct to GvHD or GvL (60 and 138, respectively) (Fig.5G). A similar trend was observed in non-responders (Fig.5H). When comparing responders with non-responders in both GvHD (Fig.5I) and GvL (Fig.5J), we found minimal overlap in the miR targetome, which was unexpected. Notably, there were only 2 putative miR targets downregulated upon miR-374b-M4 treatment in all four co-cultures types: AK4 and MRPS6. These encode mitochondrial proteins in the nucleus. Overall, these data suggest that (1) the size of the miRNA targetome defines the biological effect size, (2) GvHD and GvL share some mechanisms but also have distinct ones, and (3) miR-374b-M4 induces context-dependent metabolic and immune regulatory reprogramming in T cells during the allogeneic immune response.

### The functional state of T cells predicts response to miR-374b-M4

One key question that remains is: what distinguishes a responder from a non-responder? To address this, we compared responder and non-responder T cells either from untreated or miR-374b-M4 treated GvHD and GvL co-cultures (Fig.6). We observed a greater number of DEGs between miR-374b-M4-treated responder *versus* non-responder co-cultures (Fig.6A and 6B) (76 and 64, respectively) than between untreated co-cultures (Fig.6C and 6D) (21 and 41, respectively). This suggests that miR-374b-M4 treatment amplifies transcriptomic differences between responder and non-responder T cells. Indeed, many DEGs identified in the responder *versus* non-responder analysis of miR-374b-M4-treated co-cultures were also identified as DEGs in the treated *versus* untreated analysis of responder co-cultures. Importantly, no significant differences were found in the expression of RNAi pathway genes between responders and non-responders, indicating that the RNAi pathway is likely intact in non-responder T cells.

**Figure 6.**
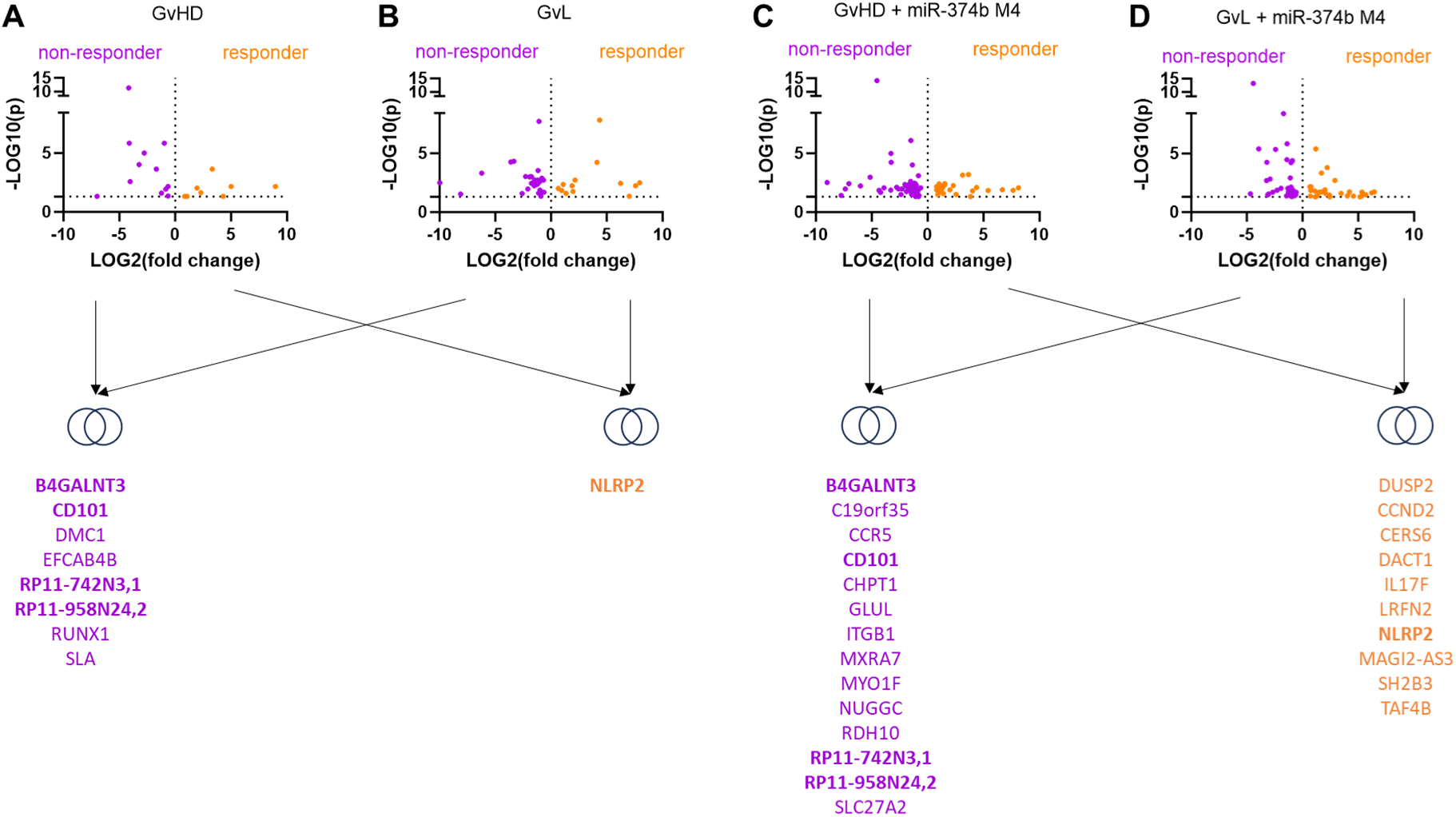
Transcriptomes define miR-374b-M4 responders. Unpaired differential expression analysis compared RNASeq data between T cells of responder and non-responder untreated GvHD (A) and GvL cultures (B), as well as miR-374b-M4-treated GvHD (C) and GvL cultures (D). On the volcano plots genes enriched in non-responder cells are depicted in purple and genes enriched in responders are depicted in orange. Genes found enriched in both GvHD and GvL responders are listed in orange and genes enriched in both GvHD and GvL non-responders are listed in purple below the Venn diagrams. Genes enriched in both untreated and miR-374b-M4-treated responders or non-responders are listed in bold. N=2

Next, we examined the overlap of DEGs enriched in non-responder GvHD co-cultures and non-responder GvL co-cultures. In untreated non-responder co-cultures, we identified 8 DEGs overlapping across GvHD and GvL, whereas in miR-374b-M4-treated co-cultures, we found 14 such DEGs (Fig.6. magenta). From this analysis, we shortlisted four genes— B4GALNT3, CD101, RP11-742N3.1, and RP11-958N24.2—that were consistently enriched across all non-responder co-cultures (Fig.6. magenta and bold). These genes were not regulated by miR374b-M4.

In untreated responder co-cultures, we found only one overlapping DEG (NLRP2) across GvHD and GvL (Fig.6. orange and bold). This gene was also found within the 11 such overlapping DEGs (Fig.6. orange) of miR-374b-M4-treated responder co-cultures. NLRP2 was not regulated by miR-374b-M4. MAGI2-AS3 was upregulated in miR-374b-M4-treated co-cultures. MAGI2-AS3 is predicted to be a miRNA sponge with 9 binding sites to the miR-374b-M4 guide strand, and is likely upregulated as a compensation mechanism to supranatural concentrations of miR-374b-M4 in responder T cells. CERS6 has been suggested as a GvHD-specific target^49^, and as a mediator of mitochondrial damage leading to STING activation and a type I interferon response^50^ – which we saw enriched in responder T cells participating in GvL.

Taken together, we identified four genes characterizing non-responders and one gene associated with responders, none of which appear to be regulated by miR-374b-M4. Among them, CD101 has been implicated in T cell exhaustion in the context of GvHD^51^. This observation aligns with our finding that non-responders were more frequently found in GvHD co-cultures with higher T cell proliferation rates (Supp. Fig.2B), a condition often linked to exhaustion. Indeed, the classical exhaustion markers HAVCR2 and LAG3 were significantly downregulated in responder miR-374b-M4 co-cultures. However, the effect size of miR-374b-M4 did not correlate with T cell proliferation in GvL assays (Supp. Fig.2C). NLRP2 is a known suppressor of NF-κB signaling^52^, a pathway suggested to be selectively involved in GvHD^53^. RP11-742N3.1 and RP11-958N24.2 are pseudogenes with uncharacterized functions. Neither contains seed matches to either strand of miR-374b, excluding a function as miRNA sponges. Notably, RP11-742N3.1, also known as RPS28P7, was highly expressed (above 150 TPM). This 374-nucleotide ribosomal pseudogene is exclusively localized in the cytoplasm^54^ and has previously been implicated in GvHD suppression^55^.

Altogether, these data suggest that the functional state of T cells predicts their responsiveness to miR-374b-M4.

## Discussion

miRNAs typically regulate multiple pathways, resulting in complex yet targeted transcriptomic rewiring in cells. This ability has sparked significant interest in miRNA replacement therapies for diseases where specific miRNAs are lost or downregulated. However, the clinical success of these therapies has been limited^5,6^, primarily because the field has yet to establish effective strategies for chemically modifying miRNA mimics in a functionally meaningful way^9^.

Notably, no studies to date have systematically explored how different chemical modification patterns of miRNA mimics affect their functional activity in disease models^9,15,17,19^.

Additionally, it remains unknown, which set of downregulated targets (*i.e.* targetome) mediate functional activity of miRNA mimics in diseases^15,17,19^. In most cases, the downregulation of a single predicted target is demonstrated, and this is assumed to account for the observed functional outcomes^26,27^—despite the well-established fact that miRNAs act through multiple targets^13^.

In this study, we used T cell responses to alloantigen as a model to investigate how chemically modified miRNA mimics drive complex transcriptomic changes. We selected three distinct miRNA sequences (miR-146a, miR-181b and miR-374b) for analysis. This approach is novel, as no previous study has simultaneously examined the chemical modification patterns of multiple miRNA sequences, which has so far precluded the development of sequence-independent, and therefore generalizable principles in the field. We found that silencing assays—whether using fully complementary reporter constructs or natural, mismatched targets—do not reliably predict the functional performance of miRNA mimics in disease models. Based on this, we propose that screening of miRNA mimic chemistries should be performed primarily in relevant functional disease models, rather than relying on standard silencing assays. As shown here, the functional effects of a miRNA mimic in disease contexts are likely mediated through the collective downregulation of potentially hundreds of genes (*i.e.* targetome).

Our findings also shed light on general principles of miRNA mimic chemistry. The relationship between chemical modification and silencing efficacy appears to be sequence-dependent, and unlike siRNAs, there may be no universally optimal chemical structure for miRNA mimics. While previous studies suggested that a fully matched sense strand impairs miRNA function^9^, we observed this only in a subset of cases, further emphasizing that such rules are context- and sequence-specific.

Interestingly, one of the most consistent chemical predictors of GvHD suppression by miRNA mimics was phosphorothioate (PS) content, though non-targeting miRNA mimics with the same PS content showed no activity. This indicates that PS-effects are unlikely to be mediated by toxicity^56^, but PS may rather amplify sequence-specific functional activity of miRNA mimics. Higher PS content in a single-stranded overhang (structures M5, M7) or in a duplexed region (structures M2, M4) mediated better functional activity of miRNA mimics equally, therefore the enhancement of functional activity by PS cannot be explained by enhanced cellular uptake (as shown in the context of single stranded PS overhangs in siRNAs)^57^. Rather the effect of PS content is likely mediated *via* enhanced protein binding^58^. We demonstrate that miRNA mimics can suppress GvHD to a degree comparable with several approved GvHD therapies in a functional disease model *in vitro*. Using two independent assays, we confirmed that miRNA mimics selectively inhibit GvHD-driving T cells, without affecting polyclonally activated T cells or impairing GvL activity. Remarkably, in some cases, we observed enhanced GvL responses of T cells upon miRNA mimic treatment, which is a highly desirable feature and contrasts with the effects of tested approved drugs, which broadly suppressed T cell activity and GvL.

These findings establish a translational path for using miRNA mimics in GvHD therapy. We propose their use in *ex vivo* treatment of adoptive cell therapies (e.g., donor lymphocyte infusion, allogeneic CAR-T cells), where they could inhibit GvHD while preserving or even enhancing GvL effects. Our data also show that the duration of miRNA mimic activity is sufficient to support this therapeutic approach. Yet, further studies are needed to establish *in vivo* proof-of-principle.

We explored the targetome of our lead candidate miRNA mimic, miR-374b-M4, in both on-target (GvHD) and off-target (GvL) contexts. We found that the magnitude of the response to miR-374b-M4 correlates with the size of the regulated targetome. The set of targeted genes differed depending on the biological context (e.g., GvHD *versus* GvL). This underlies two key points: (1) context-dependent targetomes need to be analyzed when developing miRNA-based therapies, and (2) partially different transcriptional programs underly T cell activity in GvHD and GvL. The latter suggests, that beyond TCR repertoires, transcriptional programs and functional states of T cells substantially contribute to on-*versus*-off-tumor immune responses. Our transcriptomic analyses revealed regulation of several genes and pathways previously implicated in GvHD and GvL, validating the approach. We also uncovered new insights, such as miR-374b-M4 inducing metabolic reprogramming in T cells.

We hypothesize that the response to miR-374b-M4 may be influenced by glycosylation status of CD101, a surface protein potentially glycosylated by B4GALNT3 – both enriched in non-responder T cells. We also speculate that two pseudogenes, RP11-742N3,1 and RP11-958N24,2 – both enriched in non-responder T cells – play meaningful roles in T cell biology in the context of allogeneic stimulus. These proteins and/or transcripts likely define the functional state of T cells that associate to responsiveness of miR-374b replacement therapy. In conclusion, our data support the need for functional screens for developing chemically modified miRNA mimics, and the notion that miRNA targetome profiling in disease models is a powerful tool for therapeutic target discovery.

## Methods

### miRNA mimics synthesis

Compounds were synthesized using standard solid-phase phosphoramidite chemistry on a Dr. Oligo 48 medium-throughput RNA synthesizer (Biolytic, Fremont, CA)). Standard RNA 2′-*O*-methyl and 2′-fluoro modifications were applied for improving siRNA stability (Chemgenes, Wilmington, MA and Hongene Biotech, Union City, CA). Oligonucleotides were grown on long-chain alkyl amine (LCAA) controlled pore glass (CPG) functionalized via succinyl linker with either Unylinker terminus for unconjugated oligonucleotides 500Å (Chemgenes), or with cholesterol conjugate through a tetraethylene glycol linker 500Å (Chemgenes). Oligonucleotides were cleaved and deprotected on-column with Ammonia gas (Airgas Specialty Gases). Briefly, columns were pre-wet with 100uL of water and immediately spun to remove the excess water. Columns were then placed in a reaction chamber (Biolytic) 90min at 65°C. A modified on-column ethanol precipitation protocol was used for desalting and counterion exchange. Briefly, 1mL of 0.1M sodium acetate in 80% ethanol is flushed through the column, followed by a rinse with 1mL 80% ethanol and finally after drying the excess ethanol, oligonucleotides were eluted with 600uL of water in 96 deep-well plates. The oligonucleotide fractions were quality-controlled by liquid chromatography-mass spectrometry on an Agilent 6535B QTOF LC-MS.

Oligonucleotide strands were annealed by mixing them in equal amount of substance, denaturing at 95°C, and allowing them to anneal at room temperature for at least 15 minutes. Annealing efficacy was quality-controlled using non-denaturing electrophoresis on polyacrylamide gels.

The sequences and chemical modifications patterns of all oligonucleotides used in this study are listed in Supp. Table 1.

### Cell culture

Peripheral blood mononuclear cells (PBMCs) were obtained from the buffy coats of healthy donors provided by the Transfusion Medicine Center of the University Hospital Tübingen using Ficoll (GE17-1440-02, Sigma-Aldrich) gradient centrifugation. The PBMC stocks were stored in liquid nitrogen in a freezing medium containing 90% FBS (11573397, Gibco) and 10% DMSO (APP A3672,0250, Omnilab).

T cells and dendritic cells were purified from PBMCs. 30 – 300 ×10^6^ PBMC where thawed, resuspended in 5 mL of RPMI-1640 medium (R2405, Sigma-Aldrich) with 10% FBS (11573397, Fisher Scientific) and 1% Penicillin-Streptomycin (P0781, Sigma-Aldrich), 25 mM HEPES (9157.1, Carl Roth) and 1 mM sodium pyruvate (12539059, Gibco). PBMCs were then immediately used for the purification of T cells and dendritic cells *via* negative selection. Pan T Cell Isolation Kit (130-096-535, Miltenyi Biotec) and Pan-DC Enrichment Kit (130-100-777, Miltenyi Biotec) were used according to the manufacturer’s protocols. After negative selections, cells were immediately used to set up functional disease models, as described below.

Jurkat cell line was cultured in RPMI-1640 medium supplemented with 10% FBS, 1 % P/S, 25 mM HEPES and 1 mM sodium pyruvate. Jurkat cells were maintained at a density of 0.5 to 2 ×10^6^ cells/ml by passaging three times per week.

HeLa cells were cultured in RPMI-1640 medium supplemented10% FBS, 1 % P/S, 25 mM HEPES and 1 mM sodium pyruvate. HeLa cells were passaged by detaching them using Accutase (A6964, Sigma-Aldrich) and splitting them keeping a confluence between 25% and 70%, three times a week.

All cell cultures were maintained in an incubator at 37°C with 5% CO₂.

### Cloning

An insert containing a 4x fully complementary target sequence to miR-146a, miR-374b, and miR-181a was designed and ordered through GeneArt Custom Gene Synthesis for custom DNA constructs (CLPMX5UGDE, ThermoFisher). It was cloned into psiCHECK-2 Vector (a generous gift from Dr. Anastasia Khvorova, University of Massachusetts Chan School of Medicine) in the 3’ UTR of the Renilla luciferase reporter. Plasmid and insert were digested with restriction enzymes XhoI (R0146S, New England Biolabs) and NotI (R3189S, New England Biolabs), separated by gel electrophoresis with a 1% of agarose (A8963.0250, VWR) in 1x TAE buffer. Fragments were purified from the gel with Invitrogen PureLink Quick Gel Extraction Kit (10114993, Fisher Scientific). 100 ng psiCHECK-2 and 300 ng insert were combined up to 5 µL with MilliQ water and 5 µl of Instant Sticky-end Ligase Master Mix (M0370S, New England Biolabs). 50 µl of DH5α competent cells (16506900, Fisher Scientific) where thawed, 2 µl of ligation reaction added and mixed by gently pipetting in a 1.5 centrifuge tube. The tube was incubated for 30 minutes on ice. Afterwards bacteria were transformed *via* heat-shock for 45 seconds at 42°C. A 5-minute incubation on ice followed. 950 µl of SOC medium (S1797, Sigma Aldrich) were added and kept shaking at 37°C for 1 hour and then plated on a petri dish with LB-Agar (1317GR500, Neolab) and incubated for 24h. Grown colonies were tested for the presence of the insert by Microsynth Seqlab using Sanger Sequencing with primer 5’GTCCGCAACTACAACGCCTACCTT-3’. The colony with the plasmid of interest was grown and stored in glycerol stocks by centrifuging and resuspending them in 50% glycerol (G-0650-08, ThermoFisher) and 50% milli-Q water.

### Dual-Glo®Luciferase Assay System

The Dual-Glo® Luciferase Assay System (E2920, Promega) was used to assess miRNA mimic silencing on a fully complementary target.

Plasmids were purified from DH5α using E.Z.N.A. Plasmid Mini Kit I (D6942-02, VWR). 2.5 µg of the plasmid containing 4x fully complementary sequences to miR-146a, miR-374b and miR-181a in 125 µl of OptiMEM (11520386, Fisher Scientific) were mixed with 5µl of Lipofectamine 2000 Transfection Reagent (10696343, Thermo Fisher Scientific) diluted in 120 µl of OptiMEM. The mix of plasmid, Lipofectamine 2000 and OptiMEM was incubated at room temperature for 10 minutes and added to 70% confluent HeLa cells in 3 ml of RPMI-1640 medium (R2405, Sigma-Aldrich) with 10% FBS (11573397, Fisher Scientific) and 1% Penicillin-Streptomycin (P0781, Sigma-Aldrich), 25 mM HEPES (No. 9157.1, Carl Roth) and 1 mM sodium pyruvate (12539059, Gibco). After 24 hours, transfected HeLa cells were detached using Accutase (A6964, Sigma-Aldrich) and 10 000 cells per well were plated in 100 µl of medium without phenol red (R7509, Sigma Aldrich) supplemented with 10% FBS (11573397, Fisher Scientific) and 1% Penicillin-Streptomycin (P0781, Sigma-Aldrich), 25 mM HEPES (No. 9157.1, Carl Roth) and 1 mM sodium pyruvate (12539059, Gibco). miRNA mimics were diluted in 50 µl of OptiMEM in a 1:5 dilution curve, and added to 50 µl transfected HeLa cells in each well.

After 3 days, medium was aspirated and replaced with 50 µl of fresh medium without phenol red. 50 µl of Dual-Glo® Luciferase Reagent were added without washing or preconditioning the cells, inducing lysis and acting as a substrate for firefly luciferase. Firefly luciferase luminescence was measured after 10 minutes with Tecan Infinite 200 Pro microplate reader. Then, 50 µl of Dual-Glo® Stop&Glo® Reagent were used to quench the firefly luminescence and provided a substrate for Renilla luciferase. After 10 minutes, luminescence intensity was quantified using the microplate reader. miRNA action was approximated by calculating in Renilla versus Firefly luminescence in each well.

### mRNA quantification (QuantiGene™ Singleplex assay)

Dilution curves 1:5 of miRNA mimics or miRNA non-targeting control were prepared in 50 µl OptiMEM medium (11520386, Fisher Scientific). Afterwards, 10 000 Jurkat cells were plated per well in 50 µl RPMI-1640 medium (R2405, Sigma-Aldrich) with 10% FBS (No. 11573397, Fisher Scientific) and 1% Penicillin-Streptomycin (P0781, Sigma-Aldrich), 25 mM HEPES (No. 9157.1, Carl Roth) and 1 mM sodium pyruvate (12539059, Gibco). Cells were incubated for 6 days and lysed directly in the wells by adding 50 µl of Lysis Mixture (QP0524, Thermo Fisher) with 0.1 µl of Proteinase K (QS0103, Thermo Fisher) and mixing thoroughly by pipetting (15 times). The mixture was incubated at 50-55°C for 30 minutes and used immediately or kept frozen at −80°C until use.

A probe against IRAK1 (QGS-1000, Thermo Fisher) was used to test the miR-146a mimic silencing effect, a probe against AKT1 (QGS-1000, Thermo Fisher) for miR-181a, and a probe against PTPN11 (QGS-1000, Thermo Fisher) for miR-374b. A probe against HPRT (QGS-5000, Thermo Fisher), a housekeeping gene, was used to normalize the mRNA target amount to the total mRNA amount in the cells.

### Facilitated miRNA uptake

Lipofectamine RNAiMAX Transfection Reagent (13778030, Thermo Fisher Scientific) was used to facilitate the delivery of miRNA-146a mimics into HeLa cells. miR-146a mimics were taken up in 25 µl OptiMEM at 4x the intended final concentration and were mixed with 25 µl of OptiMEM containing 0.3 µl Lipofectamine RNAiMAX. 50 µl of the supplemented RPMI-1640 medium containing 10 000 HeLa cells were added and cells were plated in 96-well tissue culture plates. Cells were incubated for 3 days and lysed directly in the wells by aspirating the medium and adding 50 µl of Lysis Mixture (QP0524, Thermo Fisher) with 100 µL of MilliQ water with 0.5 µl of Proteinase K (QS0103, Thermo Fisher) and mixing thoroughly by pipetting (15 times). The mixture was incubated at 50-55°C for 30 minutes and used immediately for mRNA quantification (see above) or kept frozen at −80°C until use.

### Duration of miRNA silencing

miR-146a fluorescent reporter plasmid was purchased from Addgene (#149718). Packaging psPAX2 (Addgene #12260), envelope pMD2.G (Addgene #12259) and corresponding transfer plasmid were transfected into HEK-293T cells at 70% confluency in 3:1:4 ratio in total amount of 20 mg to T75 flasks using 39 µl of Lipofectamine® 2000 (10696343, Fisher Scientific). After 24 h of incubation medium was changed. After 48 h virus was harvested and concentrated using 8% PEG8000 (10224963, Fischer Scientific) and 0,14 µM NaCl (10616082, Fischer Scientific). 100 µl of concentrated virus were added to 70% confluent HeLa cells in one well of 6 well plate. After 48 h medium was changed. Fluorescence of transduced cells was confirmed using microscopy (CZ-0474, Zeiss).

Transduced HeLa cells were plated at a concentration of 10 000 cells per well in 50 µl of medium without phenol red (R7509, Sigma Aldrich) supplemented with 10% FBS (11573397, Fisher Scientific) and 1% Penicillin-Streptomycin (P0781, Sigma-Aldrich), 25 mM HEPES (No. 9157.1, Carl Roth) and 1 mM sodium pyruvate (12539059, Gibco) and treated with 5 µM of miR-146a mimics. Fluorescence for mCherry and ZsGreen1 was quantified 2 times per week. Cells were passaged 1:4 twice per week.

### Graft-versus-host-disease model

PMBCs were thawed and incubated with 200 units of DNAse I in 5 ml of RPMI-1640 medium supplemented 10% FBS (11573397, Fisher Scientific) and 1% Penicillin-Streptomycin (P0781, Sigma-Aldrich), 25 mM HEPES (9157.1, Carl Roth) and 1 mM sodium pyruvate (12539059, Gibco).

Responder T cells were isolated from PBMCs using the Pan T Cell Isolation Kit (130-096-535, Miltenyi Biotec) and were labeled with CellTrace Violet Cell Proliferation Kit (15579992, Fisher Scientific): T cells in 1 ml PBS (12037539, Fisher Scientific) with 1 µM CellTrace Violet dye for 20 minutes at 37°C.

Stimulator PBMCs or dendritic cells were obtained from allogeneic donors, and were irradiated with 30 cGy using GammaCell 1000 Elite irradiator (Best Theratronics) to induce cell injury and mimic conditioning therapy prior to allogeneic stem cell transplantation. Stimulator PBMCs or dendritic cells and responder T cells were then mixed together in equal proportions (0.1×10^6^ PBMCs, 0.1×10^6^ T cells). The mixture was treated with 1, 2.5 and 5 µM of miRNA mimics or miRNA non-targeting control and an established concentration for the approved GvHD drugs that do not compromise T cell viability (7.5 µM Ruxolitinib, 5 µM Cyclosporin A, 10 µM Mycophenolat-Mofetil, 100 µM Methotrexate, 25 µM Rapamycin, 10 µM Tacrolimus, 7.5 µM Prednisolone). Cells were kept in the incubator at 37°C for 5 days. After the incubation, Zombie NIR Fixable Viability kit (423105, Biolegend) was used to stain dead cells at 1:500 dilution ratio.

Zombie NIR and CellTrace Violet staining were assessed by flow cytometry.

### Polyclonal activation of T cells

Isolated T cells were seeded in a density of 0.1×10^6^ cells per well in 100 µl of in RPMI-1640 medium (R2405, Sigma-Aldrich) with 10% FBS (11573397, Fisher Scientific) and 1% Penicillin-Streptomycin (P0781, Sigma-Aldrich), 25 mM HEPES (No. 9157.1, Carl Roth) and 1 mM sodium pyruvate (12539059, Gibco). MACSiBead Particles against CD2/CD3/CD28 (130-091-441, Miltenyi Biotec) were used to obtain activated T cells by co-incubating them in a 1:2 particle-T-cell ratio during 2-3 days at 37°C with 5% CO₂. Activated T cells were pelleted and supplemented with 10 ng/mL IL-7 (207-IL-005, Miltenyi Biotec) and 3 ng/mL IL-15 (247-ILB-005, Miltenyi Biotec).

T cells were stained with CellTrace Violet Cell Proliferation Kit (15579992, Fisher Scientific) as described above and treated with 1, 2.5 and 5 µM of miRNA mimics or non-targeting control miRNA mimics and an established concentration for the approved GvHD drugs that do not compromise T cell viability (7.5 µM Ruxolitinib, 5 µM Cyclosporin A, 10 µM Mycophenolat-Mofetil, 100 µM Methotrexate, 25 µM Rapamycin, 10 µM Tacrolimus, 7.5 µM Prednisolone). After 5 days of incubation, cells were stained with Zombie NIR viability dye and proliferation of viable T cells quantified *via* flow cytometry.

### Graft-versus-leukemia model

Stimulator dendritic cells were resuspended in 10 ml of RPMI-1640 medium (R2405, Sigma-Aldrich) with 10% FBS (No. 11573397, Fisher Scientific) and 1% Penicillin-Streptomycin (P0781, Sigma-Aldrich), 25 mM HEPES (No. 9157.1, Carl Roth) and 1 mM sodium pyruvate (12539059, Gibco) and were gamma-irradiated with 30 Gy using GammaCell 1000 Elite irradiator (Best Theratronics).

Stimulator MOLM-13 cells (acute myeloid leukemia cell line) were labelled with CellTrace Far Red Cell Proliferation Kit (C34564, Thermo Fisher) at a 1:1000 ratio.

Responder T cells were labeled with 1:1000 CellTrace Violet Cell Proliferation Kit (15579992, Fisher Scientific).

Cells were mixed by adding 1×10^5^ T cells, 1×10^5^ dendritic cells and 0.1×10^5^ MOLM-13 cells in 100 µl of medium. The mixture was treated with 5 µM of miRNA mimics or an established concentration for the approved GvHD drugs that do not compromise T cell viability (7.5 µM Ruxolitinib, 5 µM Cyclosporin A, 10 µM Mycophenolat-Mofetil, 100 µM Methotrexate, 25 µM Rapamycin, 10 µM Tacrolimus, 7.5 µM Prednisolone).

Cells were kept in the incubator at 37°C for 5 days. After the incubation, Zombie NIR Fixable Viability kit (423105, Biolegend) (1:5000) was used to stain dead cells.

### Flow cytometry

Flow cytometry was performed using a BD FACSCanto II Flow Cytometer (BD Biosciences) and BD LSRFortessa Cell Analyzer (BD Biosciences), provided by the FACS Core Facility, Berg Medical Clinic II. Flow cytometry data was analyzed using FlowJo v10.0.

Sequential gating was performed to identify live, proliferating lymphocytes in GvHD assays and polyclonally activated T cells. First, lymphocytes were gated based on their forward and side scatter characteristics, selecting the population with intermediate FSC-A and low SSC-A to exclude debris and granulocytes. Next, doublets and aggregates were excluded by plotting forward scatter area against height (FSC-A vs. FSC-H) and gating along the diagonal representing single cells. Viability staining was then used to exclude dead cells. Cells negative for the viability dye Zombie NIR were gated as live cells, while positively stained cells were excluded. Proliferating cells were identified by progressive dilution of the CellTraceViolet dye, with undivided cells exhibiting the highest fluorescence intensity. Successive peaks representing each round of cell division were quantified using histogram plots of dye fluorescence. Gating thresholds were established using unstained and freshly stained controls as well as resting T cells to define the undivided population.

In GvL assays both cells were first assigned to either T cells or MOLM-13 based on positivity of CellTraceViolet (T cells) or CellTraceRed (MOLM-13). Then dead and live cells were quantified in each group based on Zombie NIR staining. Proliferating T cells were quantitated similarly than in GvHD assays. Resting T cells, MOLM-13 only cultures as well as unstained controls were used to set the gates.

The gating strategy used in this study is depicted in Supp. Fig.3-4.

### RNA-Sequencing

For GvHD samples, isolated T cells were mixed together in equal proportions with isolated dendritic cells from allogeneic donors (0.1×10^6^ T cells, 0.1×10^6^ gamma-irradiated dendritic cells in 100 µl per well, repeated for 15 wells). Cell mixtures were treated with 5 µM of miR-374b M4 or left untreated.

For GvL samples, 0.1×10^6^ T cells, 0.1×10^6^ allogeneic, gamma-irradiated dendritic cells and 0.01×10^6^ MOLM-13 cells were mixed in 100 µl of medium per well, repeated for 15 wells. The mixture was treated with 5 µM of miR-374b M4 or left untreated.

Each GvHD and GvL reaction was set up simultaneously in 14 wells in a 96 well plate. One well was used for flow cytometry analysis in order to classify the samples as responders or non-responders. The rest of the wells were pulled together and T cells were re-isolated from co-cultures using the Pan T Cell Isolation Kit (130-096-535, Miltenyi Biotec). Afterwards, RNA was isolated using the Monarch Total RNA Miniprep Kit (T2010S, New England Biolabs). Quality of the samples were measured with 2100 Bioanalyzer Instrument (Agilent).

Library preparation and sequencing were performed by the NGS Core Facility (NEBNext Ultra II Directional RNA Library Prep Kit for Illumina (E7760S), NovaSeq 6000 System (20040719, Illumina)). The Center of Quantitative Biology performed RNA-seq primary and secondary data analysis: mapping to the human genome, gene assignment, read assignment, raw and normalized gene counts, differential expression analysis using DESeq2, mapping of seed matches to 3’UTRs of detected genes.

Gene ontology analysis was performed using DAVID^59,60^.

### Data Visualization

Data was visualized using Prism version 10.1.1.

## Acknowledgement

We thank Anastasia Khvorova for making oligonucleotide synthesis infrastructure available for this project. We thank the FACS Core Facility, University Hospital Tübingen for assistance with flow cytometric measurements. We acknowledge the NGS Core Facility and the Center of Quantitative Biology of the University of Tübingen for performing RNASeq and primary data analysis.

This work was supported by the German Cancer Aid [70113948 to R.A.H.]; and the Faculty of Medicine, University of Tübingen [473-0-0 to R.A.H., 2652-0-0 to R.A.H.]. R.A.H. was further supported by the MINT-Clinician Scientist program of the Medical Faculty Tübingen (DFG, German Research Foundation, 493665037), and the Deutsche Gesellschaft für Innere Medizin Clinician Scientist Program. This work was partly funded by the Deutsche Forschungsgemeinschaft (DFG, German Research Foundation, 467578951), NIH grant S10OD020012 for the Mid-Scale RNA Synthesis, Purification, and Quality Control System, and NIH grant S10OD036329 for the High-throughput Oligonucleotide Production System.

## Author Contributions

Conceptualization X.S., R.A.H. Methodology X.S., A.K., T.R., R.A.H., Investigation X.S., T.R., M.S., S.J., D.A.C., D.E., Writing – Original Draft R.A.H. Visualization X.S., R.A.H. Supervision R.A.H. Project Administration R.A.H. Funding Acquisition R.A.H.

