## Supplementary Material for "miRNA Mimic Optimization: Chemical Structure – Activity – Targetome Relationship to Engineer Selective Anti-Tumor Immunity in T cells"

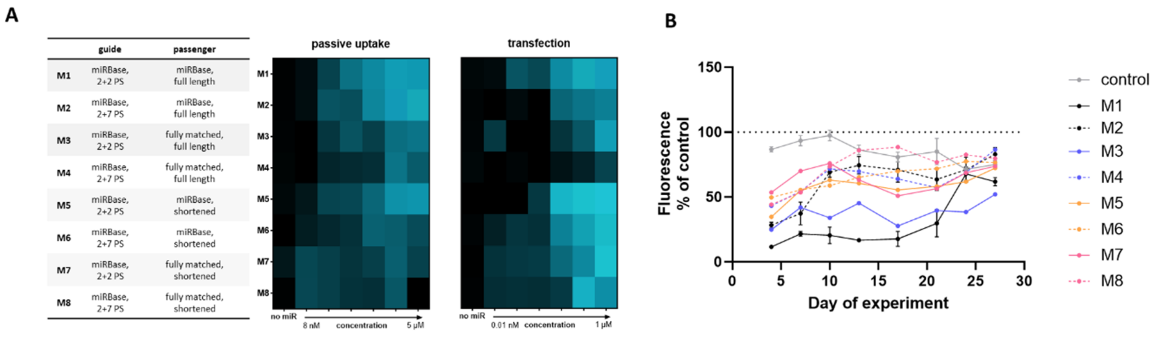


**Supplementary Figure 1.** (A) *IRAK1* mRNA silencing was quantified in Jurkat cells after co-incubation with (passive uptake) or in HeLa cells after RNAiMAX-mediated transfection with (transfection) miR-146a mimics at varying concentrations as depicted on the x axis. *IRAK1* expression was analyzed using QuantiGene SinglePlex assay and *HPRT* as housekeeping gene. Gene expression data was the normalized to untreated samples and colorcoded with black representing no silencing and blue representing maximal silencing. N=8-9. (B) HeLa cells were transduced with a fluorescent miR-146a sensor and treated with miR-146a at 5 µM. mCherry (miR-146a target) and ZsGreen1 (transduction efficiency control) was quantified using a fluorescent plate reader twice a week for 27 days. Data was normalized to untreated samples (dashed line). N=6, mean ± SEM

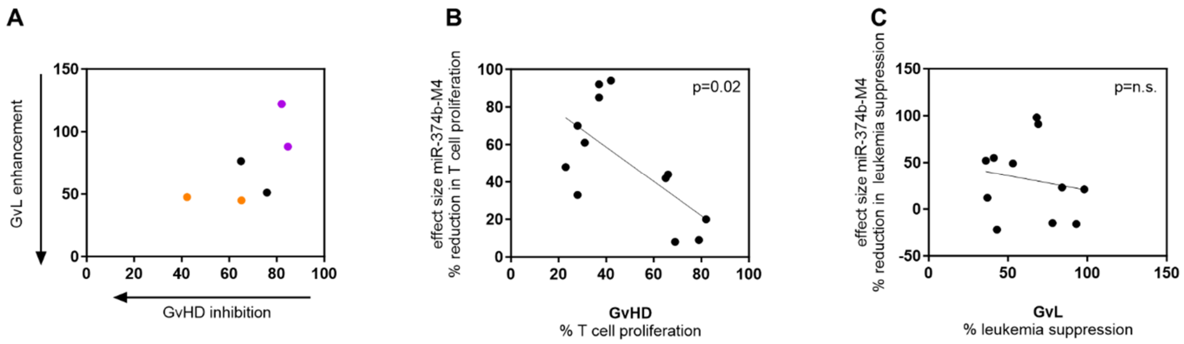


**Supplementary Figure 2.** (A) GvHD and GvL assays were performed with the same T cells and same allogeneic dendritic cells in miR-374b-M4-treated and untreated samples. On the x axis T cell proliferation in GvHD assays normalized to untreated samples is depicted, with lower numbers representing more potent GvHD suppression. On the y axis the number or viable leukemia cells in GvL assays normalized to untreated samples are depicted, with lower number representing more potent enhancement of the GvL effect. We selected the co-cultures in purple as non-responder for both assays, and the co-cultures in orange as responders for both assays for downstream RNASeq analysis. (B) All GvHD experiments performed with miR-374b-M4 are visualized: The x axis shows the percent of proliferating T cell among all T cells in untreated GvHD co-cultures with higher numbers suggesting more alloreactivity in any given experiment. The y axis represents the precent reduction in T cells proliferation induced by miR-374b-M4 compared to untreated GvHD co-cultures, with lower numbers indicating more potent GvHD suppression. (C) All GvL experiments performed with miR-374b-M4 are visualized: The x axis shows the percent of leukemia suppression in untreated GvL co-cultures compared to leukemia-only-cultures with lower numbers suggesting more potent GvL effect in any given experiment. The y axis represents the precent reduction in leukemia suppression induced by miR-374b-M4 compared to untreated GvL co-cultures, with positive numbers indicating GvL enhancement and negative numbers indicating GvL suppression.

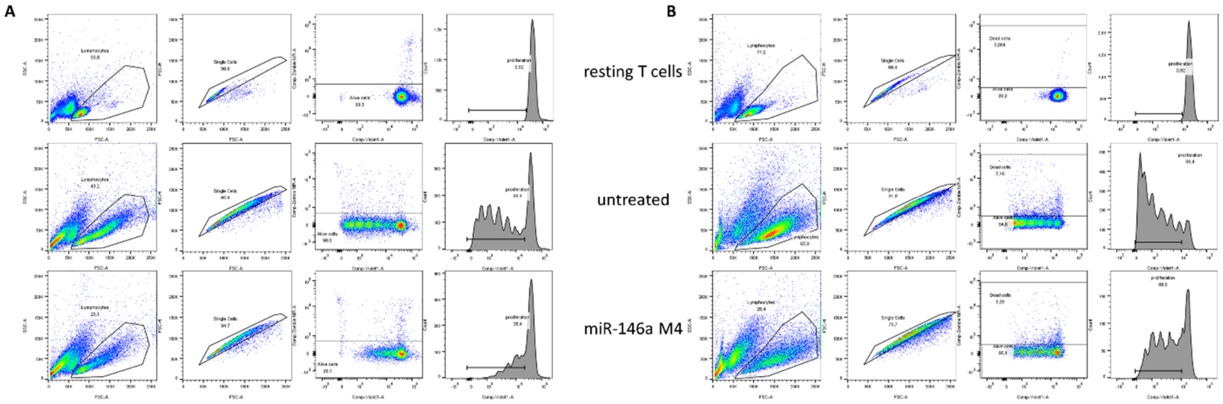


**Supplementary Figure 3.** Gating strategy in (A) GvHD assays and in (B) polyclonally activated T cells to quantitate the proportion of proliferating cells among all T cells.

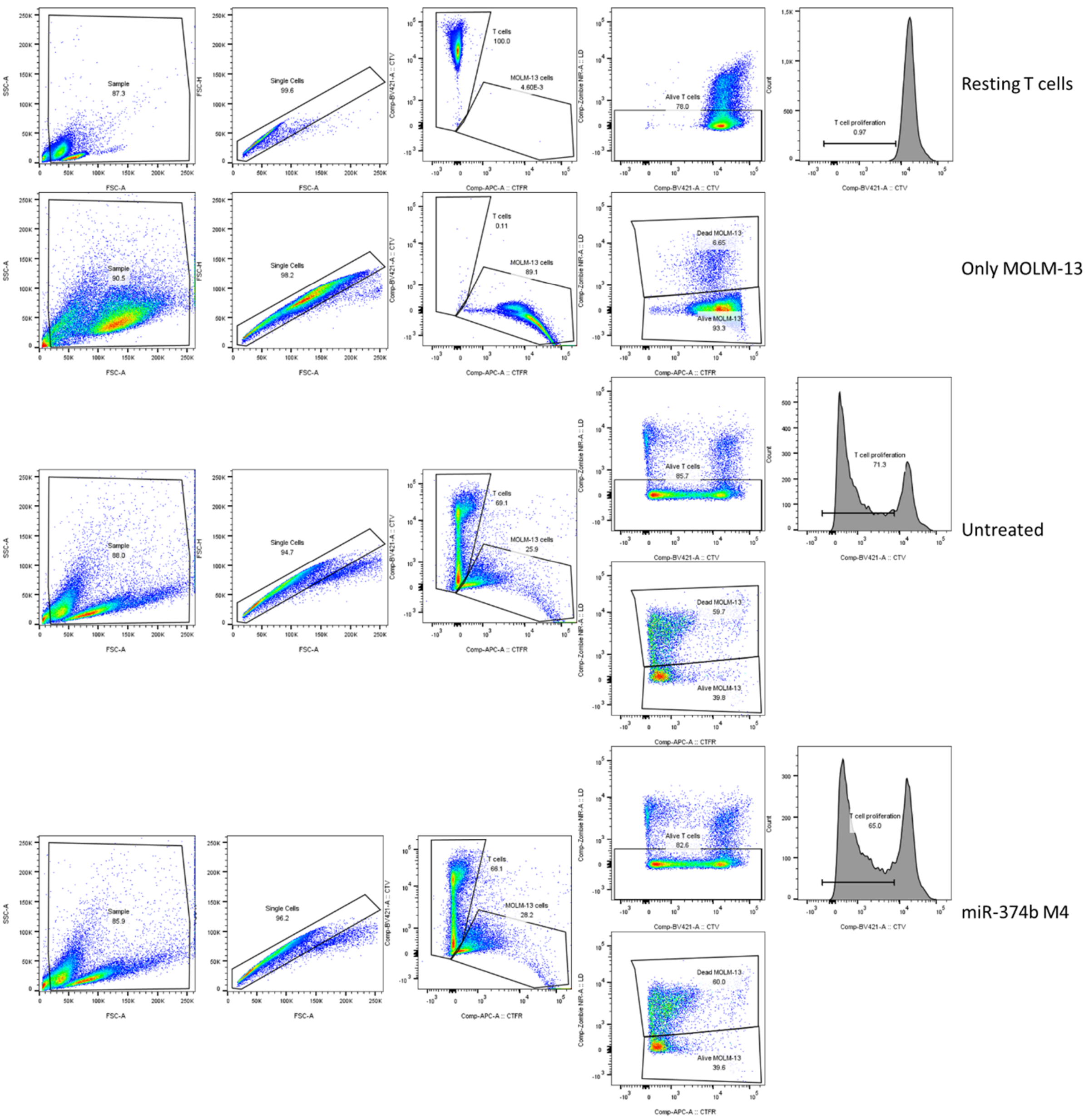


**Supplementary Figure 4.** Gating strategy in GvL assays to quantitate viable and dead leukemia (MOLM-13) cells as well as the proportion of proliferating cells among all viable T cells.

| | | |
|---|---|---|
| **miR-146a** | **Guide**<br>2+2 PS | P(mU)#(fG)#(mA)(fG)(mA)(fA)(mC)(fU)(mG)(fA)(mA)(fU)(mU)(fC)(mC)(fA)(mU)(fG)(mG)(fG)#(mU)#(fU) |
| | **Guide**<br>2+7 PS | P(mU)#(fG)#(mA)(fG)(mA)(fA)(mC)(fU)(mG)(fA)(mA)(fU)(mU)(fC)(mC)#(fA)#(mU)#(fG)#(mG)#(fG)#(mU)#(fU) |
| | **Passenger**<br>miRbase<br>full length | (mC)#(fC)#(mU)(fC)(mU)(fG)(mA)(fA)(mA)(fU)(mU)(fC)(mA)(fG)(mU)(fU)(mC)(fU)(mU)(fC)#(mA)#(fG)-TegChol |
| | **Passenger**<br>Fully matched, full length | (mA)#(fA)#(mC)(fC)(mC)(fA)(mU)(fG)(mG)(fA)(mA)(fU)(mU)(fC)(mA)(fG)(mU)(fU)(mC)(fU)#(mC)#(fA)-TegChol |
| | **Passenger**<br>miRBase, shortened | (mC)#(fC)#(mU)(fC)(mU)(fG)(mA)(fA)(mA)(fU)(mU)(fC)(mA)(fG)(mU)#(fU)#(mC)-TegChol |
| | **Passenger** fully matched, shortened | (mA)#(fA)#(mC)(fC)(mC)(fA)(mU)(fG)(mG)(fA)(mA)(fU)(mU)(fC)(mA)#(fG)#(mU)-TegChol |
| **miR-181a** | **Guide**<br>2+2 PS | P(mA)#(fA)#(mC)(fA)(mU)(fU)(mC)(fA)(mA)(fC)(mG)(fC)(mU)(fG)(mU)(fC)(mG)(fG)(mU)(fG)(mA)#(fG)#(mU) |
| | **Guide**<br>2+7 PS | P(mA)#(fA)#(mC)(fA)(mU)(fU)(mC)(fA)(mA)(fC)(mG)(fC)(mU)(fG)(mU)(fC)#(mG)#(fG)#(mU)#(fG)#(mA)#(fG)#(mU) |
| | **Passenger**<br>miRbase<br>full length | (mA)#(fC)#(mC)(fA)(mC)(fU)(mG)(fA)(mC)(fC)(mG)(fU)(mU)(fG)(mA)(fC)(mU)(fG)(mU)(fA)#(mC)#(fC)-TegChol |
| | **Passenger**<br>Fully matched, full length | (mA)#(fC)#(mU)(fC)(mA)(fC)(mC)(fG)(mA)(fC)(mA)(fG)(mC)(fG)(mU)(fU)(mG)(fA)(mA)(fU)(mG)#(fU)#(mU)-TegChol |
| | **Passenger**<br>miRBase, shortened | (fU)#(mG)#(fA)(mC)(fC)(mG)(fU)(mU)(fG)(mA)(fC)(mU)(fG)(mU)(fA)#(mC)#(fC)-TegChol |
| | **Passenger** fully matched, shortened | (fC)#(mC)#(fG)(mA)(fC)(mA)(fG)(mC)(fG)(mU)(fU)(mG)(fA)(mA)(fU)(mG)#(fU)#(mU)-TegChol |
| **miR-374b** | **Guide**<br>2+2 PS | P(mA)#(fU)#(mA)(fU)(mA)(fA)(mU)(fA)(mC)(fA)(mA)(fC)(mC)(fU)(mG)(fC)(mU)(fA)(mA)(fG)#(mU)#(fG) |
| | **Guide**<br>2+7 PS | P(mA)#(fU)#(mA)(fU)(mA)(fA)(mU)(fA)(mC)(fA)(mA)(fC)(mC)(fU)(mG)#(fC)#(mU)#(fA)#(mA)#(fG)#(mU)#(fG) |
| | **Passenger**<br>miRbase<br>full length | (mC)#(fU)#(mU)(fA)(mG)(fC)(mA)(fG)(mG)(fU)(mU)(fG)(mU)(fA)(mU)(fU)(mA)(fU)(mC)(fA)#(mU)#(fU)-TegChol |
| | **Passenger**<br>Fully matched, full length | (mC)#(fA)#(mC)(fU)(mU)(fA)(mG)(fC)(mA)(fG)(mG)(fU)(mU)(fG)(mU)(fA)(mU)(fU)(mA)(fU)#(mA)#(fU)-TegChol |
| | **Passenger**<br>miRBase, shortened | (fC)#(mA)#(fG)(mG)(fU)(mU)(fG)(mU)(fA)(mU)(fU)(mA)(fU)(mC)(fA)#(mU)#(fU)-TegChol |
| | **Passenger** fully matched, shortened | (fA)#(mG)#(fC)(mA)(fG)(mG)(fU)(mU)(fG)(mU)(fA)(mU)(fU)(mA)(fU)#(mA)#(fU)-TegChol |
| **non-targeting control** | **Guide**<br>2+2 PS | P(mU)#(fU)#(mA)(fA)(mU)(fC)(mU)(fC)(mU)(fU)(mU)(fA)(mC)(fU)(mG)(fA)(mU)(fA)(mU)#(fA)#(mU) |
| | **Guide**<br>2+7 PS | P(mU)#(fU)#(mA)(fA)(mU)(fC)(mU)(fC)(mU)(fU)(mU)(fA)(mC)(fU)#(mG)#(fA)#(mU)#(fA)#(mU)#(fA)#(mU) |
| | **Passenger**<br>miRbase<br>full length | (mA)#(fA)#(mU)(fU)(mA)(fG)(mA)(fG)(mA)(fA)(mA)(fU)(mG)(fA)(mC)(fU)(mA)(fU)(mA)#(fU)#(mA)-TegChol |
| | **Passenger**<br>Fully matched, full length | (mA)#(fU)#(mA)(fU)(mA)(fU)(mC)(fA)(mG)(fU)(mA)(fA)(mA)(fG)(mA)(fG)(mA)(fU)(mU)#(fA)#(mA)-TegChol |
| | **Passenger**<br>miRBase, shortened | (mA)#(fA)#(mU)(fU)(mA)(fG)(mA)(fG)(mA)(fA)(mA)(fU)(mG)(fA)#(mC)#(fU)-TegChol |
| | **Passenger** fully matched, shortened | (mA)#(fU)#(mA)(fU)(mA)(fU)(mC)(fA)(mG)(fU)(mA)(fA)(mA)(fG)#(mA)#(fG)-TegChol |

**Supplementary Table 1.** Sequences and chemical modification patterns used in this study. P: 5'-phosphate, #:Phosphorothioate linkage, m: 2'-O-methyl, f: 2'-fluoro, TegChol: Triethyl-glycol-linker conjugated cholesterol
